# ABHD2 activity is not required for the non-genomic action of progesterone on human sperm

**DOI:** 10.1101/2024.12.17.628646

**Authors:** Madison Edwards, Alexandra Amaral, Eve M. Carter, Oliver Arnolds, Karen Vester, Anna Thrun, Edvard Wigren, Evert Homan, Pauline Ribera, Kirsty Bentley, Martin Haraldsson, Nmesoma Theo-Emegano, Peter Loppnau, Magdalena M. Szewczyk, Michelle A. Cao, Dalia Barsyte-Lovejoy, Nicole Dittmar, Anika Hans, Mandy Weber, Jens Münchow, W. Felix Zhu, Louisa Temme, Christoph Brenker, Timo Strünker, Michael Sundström, Matthew H. Todd, Aled M. Edwards, Ralf Lesche, Opher Gileadi, Claudia Tredup

## Abstract

**Study question:** Is the hydrolase ABHD2 required for progesterone-induced Ca^2+^ influx via CatSper and the resulting motility responses in human sperm?

**Summary answer:** Progesterone-induced Ca^2+^ influx via CatSper and the resulting motility responses in human sperm do not require ABHD2 activity.

**What is known already:** Sperm motility is tightly regulated by signalling pathways that are activated as sperm ascend the female reproductive tract, including progesterone triggering Ca^2+^ influx via the CatSper channel and inducing hyperactivated motility needed for fertilization. This process is thought to involve ABHD2, which may hydrolyse the endogenous CatSper inhibitor 2-arachidonoylglycerol (2-AG), thereby relieving inhibition and enabling calcium entry into the flagellum.

**Study design, size, duration:** Potent small molecule inhibitors of ABHD2 activity were synthesized, characterized, and used as tools to scrutinize the role of ABHD2 in activation of CatSper and regulation of sperm motility.

**Participants/materials, setting, methods:** Derivatives of published ABHD2 inhibitors were optimised for *in vitro* potency and cellular activity and subsequently tested in human sperm motility and Ca^2+^ influx assays.

**Main results and the role of chance:** Progesterone does not activate ABHD2 in vitro. In addition, inhibition of ABHD2 in human sperm has no effect on progesterone-induced Ca^2+^ influx through CatSper nor on basal or progesterone-induced hyperactivated motility. This demonstrates that ABHD2 activity is, in fact, not required for the non-genomic action of progesterone on human sperm.

**Large scale data:** none

**Limitations, reasons for caution:** We examined the effects of inhibition of the enzymatic activity of ABHD2. We cannot exclude that ABHD2 functions as a part of a larger multiprotein complex, in which it may play a structural role independent of its hydrolase activity.

**Wider implications of the findings:** This study presents conclusive evidence that ABHD2 does not bind progesterone and that its hydrolase activity is not required for progesterone activation of CatSper and resulting changes in motility of human sperm. These results highlight the need for further research to elucidate the mechanism underlying the non-genomic action of progesterone on human sperm.

**Study funding/competing interest(s):** This publication is based on research funded by the Gates Foundation. The findings and conclusions contained within are those of the authors and do not necessarily reflect positions or policies of the Gates Foundation. LT, CB, and TS were supported by the Deutsche Forschungsgemeinschaft (DFG, German Research Foundation) - project numbers 329621271 (CRU326; CB, TS), and 404595355 (Research Training Group “Chemical biology of ion channels (Chembion)’; LT, TS). This work was funded by the German Federal Ministry of Education and Research (BMBF) within the framework of Contraception Research, grant number 01GR2501A and 01GR2502A.

A.A., K.V., A.T., N.D., A.H., M.W. and R.L. are employees of Nuvisan ICB GMBH, Berlin, Germany. Nuvisan is recipient of a Gates Foundation grant. A.A is associate editor of Human Reproduction Open.

## Introduction

During their journey through the female reproductive tract, human sperm are exposed to progesterone, a steroid hormone released by the cumulus cells surrounding the egg. Progesterone triggers a rapid increase in sperm intracellular calcium (Ca^2+^) levels via a non-genomic mechanism, initiating several physiological responses critical for successful fertilisation (Blackmore et al., 1990, Harper et al., 2004, Luo et al., 2019, Publicover et al., 2007, Young et al., 2024). One such response is sperm hyperactivation, a distinct motility pattern characterized by high amplitude, asymmetrical flagellar bending (Suarez, 2008).

It is well established that progesterone activates the sperm-specific Ca^2+^ channel CatSper (Lishko and Kirichok, 2010, Strunker et al., 2011), although the underlying molecular mechanism has remained elusive. It has been proposed that the serine hydrolase ABHD2 (α/β hydrolase domain-containing protein 2) functions as a membrane progesterone receptor, mediating CatSper activation (Miller et al., 2016). According to this model, ABHD2 catalyses the hydrolysis of 2-arachidonoylglycerol (2-AG), an endocannabinoid that is abundant in sperm membranes and may act as an endogenous inhibitor of CatSper. Upon progesterone binding, ABHD2 would become enzymatically active, degrading 2-AG and thereby relieving its inhibitory effect on CatSper. The resulting activation of CatSper channels would promote Ca^2+^ influx and sperm hyperactivation.

Given its proposed role in the control of sperm function, ABHD2 has emerged as a promising target for the development of non-hormonal contraceptives. Supporting this idea, decreased ABHD2 mRNA levels and frame-shift mutations have been identified in infertile men with oligoasthenozoospermia, a condition characterised by reduced sperm count and impaired sperm motility (Jiang et al., 2021).

To better understand the physiological role of ABHD2 in sperm and to assess its potential as a contraceptive target, the development of selective small-molecule inhibitors is essential. Miller et al. employed the broad-spectrum serine hydrolase inhibitor methyl arachidonyl fluorophosphonate (MAFP), which was reported to suppress progesterone-induced CatSper activation, Ca^2+^ influx and sperm hyperactivation (Miller et al., 2016). In a follow-up study, various steroids and steroid-like compounds were evaluated for their impact on CatSper-mediated Ca^2+^ signalling, identifying the plant triterpenoids pristimerin and lupeol as inhibitors of progesterone-induced Ca^2+^ influx and hyperactivation (Mannowetz et al., 2017). However, these findings were later challenged by independent studies reporting that pristimerin and lupeol do not interfere with progesterone-induced CatSper activation (Brenker et al., 2018, Rehfeld, 2020, Rehfeld and Marcus Pedersen, 2022). Notably, whether these plant triterpenoids inhibit ABHD2 enzymatic activity remains to be elucidated. To date, the only selective ABHD2 inhibitor reported is *Compound 183* (Compound **1** in the present work), which was identified using activity-based protein profiling (ABPP; (Baggelaar et al., 2019)). This compound was reported to block progesterone-induced Ca^2+^ influx in mouse sperm (Baggelaar, et al., 2019); It is, however, well established that mouse CatSper is not activated by progesterone (Lishko et al., 2011), rendering results on the action of the compound on mouse sperm difficult to interpret. In human sperm, the compound seems to have no effect on basal hyperactivation (Gruber et al., 2024); whether it impacts progesterone-induced Ca^2+^ influx and hyperactivation is not known.

This study aimed to determine whether ABHD2 activity is required for the non-genomic action of progesterone in human sperm, specifically its ability to trigger CatSper-mediated Ca^2+^ influx and hyperactivation. Our findings demonstrate that pharmacological inhibition of ABHD2 does not affect progesterone-induced Ca^2+^ signalling or hyperactivation. Furthermore, neither progesterone nor other steroids or plant-derived triterpenoids bind to ABHD2 or modulate its enzymatic activity.

## Materials and methods

### Chemicals

All reagents were from Merck KGaA (Darmstadt, Germany), unless otherwise stated.

### Biological material

Fresh semen samples were obtained from healthy, normozoospermic donors (as defined by World Health Organisation criteria; WHO, 2021) who provided prior written informed consent and observed 3-5 days of sexual abstinence prior to collection. For sperm Ca^2+^ experiments, samples were collected under approvals from the Ethical Committees of the Medical Association Westfalen-Lippe and the Medical Faculty of the University of Münster, Germany (approval ID: 4INie, 2021-402-f-S). For sperm hyperactivation experiments, samples were sourced from CRS Clinical Research Services Berlin GmbH (Germany), which operates in compliance with all applicable laws, regulations, and relevant governmental and regulatory standards for human sample donation. In both cohorts, semen was produced by masturbation, after 3-5 days abstinence, and ejaculated into sterile plastic containers.

### Expression and purification of recombinant proteins

#### Full-length ABHD2 (ABHD2^FL^)

The full-length ABHD2 sequence was codon-optimised for insect cells and cloned into a baculovirus vector with a C-terminal TEV site followed by an eGFP-8xHis tag. For protein expression, SF9 cells were infected with a multiplicity of infection of 0.1 of the third virus generation and the cell pellet was harvested 48 h after infection by centrifugation. For cell lysis, the pellet was resuspended in solubilisation buffer (50 mM Tris-HCl pH 7.5 at room temperature, 500 mM NaCl, 10 mM imidazole, 2 mM EDTA, 0.5 mM DTT, 1 % (w/v) DDM/0.1 % CHS (w/v) solution (Anatrace, Maumee, OH, USA); 0.1 μg/ml benzonase) and incubated for 1 h at 4 °C. Solubilisation was facilitated by 7-10 strokes with a glass Dounce homogeniser. After centrifugation at 35,000 × g for 1 h at 4 °C, the supernatant was harvested, filtered through a 125 µM nylon mesh and diluted 1.7-fold in wash buffer (50 mM Tris-HCl pH 7.5 at room temperature, 500 mM NaCl, 40 mM imidazole, 1 mM DTT, 0.1 % (w/v) DDM and 0.01 % (w/v) CHS). The diluted supernatant was loaded onto a His trap FF crude affinity column, washed and eluted via a continuous gradient to elution buffer (50 mM Tris-HCl pH 7.5 at room temperature, 300 mM NaCl, 400 mM imidazole, 1 mM DTT, 0.1 % (w/v) DDM and 0.01 % (w/v) CHS). The C-terminal eGFP-8xHis tag was cleaved overnight upon addition of TEV protease and incubation during dialysis into dialysis buffer (50 mM Tris-HCl pH 7.5 at room temperature, 300 mM NaCl, 10 mM imidazole, 2 mM DTT, 0.1 % (w/v) DDM and 0.01 % (w/v) CHS). The protein was loaded again onto a HisTrap™ FF crude column equilibrated in wash buffer and the flow-through of the re-chromatography was collected, containing the cleaved ABHD2 protein. The protein was concentrated with a 30 kDa cutoff filter (Amicon GmbH, Borken, Germany) and loaded onto a S200 10/300 column (GE HealthCare GmbH, Düsseldorf, Germany) equilibrated in SEC buffer (50 mM Tris-HCl pH 7.5 at room temperature, 300 mM NaCl, 1 mM DTT, 0.1 % (w/v) DDM and 0.01 % (w/v) CHS). Fractions of the chromatogram peak containing the ABHD2 protein were concentrated to 2 mg/ml as determined via nanodrop measurements and flash-frozen in liquid nitrogen. For quality control analysis, protein bands of the SDS-PAGE from the purification were cut, digested and subjected to LC-MS/MS analysis.

#### Truncated ABHD2 (ABHD2^L33–E425^)

The gene for human ABHD2 protein (Uniprot ID P08910) was introduced through Ligation Independent Cloning with In-Fusion enzyme (Takara Bio USA, Inc, San Jose, CA, USA) into several vectors for expression tests. After testing various expression constructs in *E. coli*, SF9 cells, and mammalian cells, we were able to obtain expression for ABHD2 amino acids L33 - E425, in which a predicted transmembrane domain (amino acids 1-32) is removed. This construct was expressed in the *E. coli* expression vector pET28-FLAG which was derived from pET28a(+). This vector confers kanamycin resistance and expresses an N-terminal His6-FLAG sequence under T7/Lac regulation. The resulting plasmids were propagated in *E. coli* DH5α cells, and the sequence verified. For expression, the plasmids were transformed into chemically competent BL21(DE3)-R3-pRARE2 *E. coli* cells and propagated in medium containing kanamycin (50 µg/ml) and chloramphenicol (34 µg/ml).

*E. coli* cells were grown in Terrific Broth medium with 0.8 % glycerol, chloramphenicol and kanamycin, in UltraYield flasks or the LEX bioreactor system (Harbinger Biotechnology and Engineering Corp., ON, Canada) and each flask was inoculated with overnight culture (0.01:1 overnight culture:media). The cells were incubated at 37 °C until the OD600 reached□∼□1.0; this step generally took 3-4 h. The cultures were then cooled for 20 min by reducing the temperature to 15 °C. The expression of each protein was induced with the addition of IPTG to a final concentration of 0.3 mM. The induction was carried out for 14-18 h at 15 °C.

After overnight induction, the cells were harvested by centrifuging at 6,250 × g for 15 min. Cell pellets were resuspended in Sonication Buffer (50 mM Tris pH 8, 500 mM NaCl, 10 % (v/v) Glycerol, 1% (v/v) Triton X-100, 2 mM TCEP). Serine protease inhibitors were not added, as these could inhibit ABHD2. Resuspended cells were sonicated for 7 min (total sonication time) in intervals of 8 s on, 30 s off, followed by a 10 min incubation with 3 µg benzonase per 100ml resuspended cells. Cell debris was removed by centrifugation at 36,000 × g for 50 min. The clarified lysate was incubated with nickel-sepharose beads (5 ml of beads per 10 l of cells), then the beads were washed with 20 CV Lysis buffer (50 mM Tris pH 8, 500 mM NaCl, 10 mM Imidazole, 10% (v/v) Glycerol, 2 mM TCEP), 32 CV Wash Buffer (50 mM Tris pH 8, 1 M NaCl, 30 mM Imidazole, 2 mM TCEP) and 4 CV Lysis buffer to lower the salt concentration. Samples were eluted in Elution Buffer (50 mM Tris pH 8, 500 mM NaCl, 250 mM Imidazole, 10% (v/v) Glycerol, 2 mM TCEP). Purification was carried out either by gravity flow or on an AKTAxpress purification machine using HisTrap-excel columns.

Elution fractions were pooled, concentrated to 5 ml, filtered via a 0.45 µm syringe filter, and loaded on a HiLoad 16/600 Superdex 75 pg. The protein eluted as a tetramer, and pure elution fractions were concentrated in 20 mM HEPES pH 8, 500 mM NaCl, 10 % (v/v) glycerol, 1 mM TCEP, snap frozen in liquid nitrogen, and stored at -80 °C. For quality control analysis, purified protein was analysed by LC-MS/MS.

#### ABHD10 and ABHD11

Constructs encoding amino acids T53 – N306 of human ABHD10 and residues V34 – V315 of human ABHD11 were cloned in vector pNIC28-Bsa4^40^, which adds a N-terminal His6 tag followed by a TEV protease cleavage site. Both proteins were expressed and purified from *E. coli* cells as described for *ABHD2*^*L33–E425*^, except that Triton X-100 was not included in the lysis buffer.

### Enzymatic activity assays

#### ABHD2^FL^

Enzymatic activity of ABHD2^FL^ was measured using the fluorogenic substrates 7-hydroxycoumarinyl arachidonate (7-HCA) (Sigma-Aldrich #U0383) and resorufin butyrate (RB) (Sigma-Aldrich #83637). Assays were performed in low-volume black 384-well plates (Greiner BioOne GmbH, Frickenhausen, Germany, #784076) with a final volume of 5 µl per well (2 µl enzyme or buffer plus 3 µl substrate) in an aqueous assay buffer comprising 50 mM HEPES-KOH (pH 7.5), 150 mM NaCl, and 0.01% (v/v) Triton X-100. Three experiment types were conducted: enzyme titrations, Michaelis–Menten kinetics and compound testing, with buffer-only wells included as blanks.

For the 7-HCA assay, ABHD2^FL^ was prepared as seven 2-fold serial dilutions, starting at 200 nM to yield final concentrations of 3.125, 6.25, 12.5, 25, 50, 100, and 200 nM. Reactions were initiated by the addition of 3 µl 7-HCA at a final concentration of 5 µM and incubated for 75 min at room temperature. Fluorescence from product formation was measured using an Infinite M1000 PRO plate reader (Tecan, Männedorf, Switzerland) with excitation at 335 nm and emission at 450 nm.

For Michaelis–Menten kinetics, enzymatic activity was monitored fluorometrically over time to determine initial velocities across a range of substrate concentrations. Reactions were initiated by incubating 6 nM ABHD2^FL^ with varying concentrations of 7-HCA (µM): 1.5, 3.1, 6.25, 12.5, 25, 50 and 100. Fluorescence was recorded every 0.5 min throughout a 60 min incubation. The slope of the linear segment of each substrate concentration was obtained by linear regression and initial velocities were plotted as a function of substrate concentration. Michaelis constant (Km) was determined by nonlinear regression to the Michaelis-Mention equation using GraphPad Prism (Version 10, GraphPad Software).

For compound testing in this assay, 15 nM of 50 nM ABHD2^FL^ (depending on enzyme batch) was used, and samples were pre-incubated with 50 nl or DMSO or of 50 nl of 100-fold concentrated test compounds in DMSO for 15 min at room temperature prior to addition of 7-HCA substrate at a final concentration of 5 µM. Fluorescence measurement was performed after a 40 min incubation at room temperature.

For the RB assay, ABHD2^FL^ was prepared as seven 2-fold serial dilutions, starting at 100 nM to yield final enzyme concentrations of 1.56, 3.1, 6.25, 12.5, 25, 50, and 100 nM. Reactions were initiated by adding of 3 µl RB at a final concentration of 10 µM and incubated for 60 min at room temperature. Fluorescence was recorded using a PHERAstar® *FSX* plate reader (BMG LABTECH, Ortenberg, Germany), with excitation at 540 nm and emission at 590 nm.

For Michaelis–Menten kinetics, enzymatic activity was monitored fluorometrically over time to determine initial velocities across a range of substrate concentrations. Reactions were initiated by incubating 25 nM ABHD2^FL^ with varying concentrations of RB (µM): 7.8, 15.6, 31.3, 62.5, 125 and 250. Fluorescence was recorded every 0.5 min throughout a 60 min incubation. The slope of the linear segment of each substrate concentration was obtained by linear regression and initial velocities were plotted as a function of substrate concentration. Michaelis constant (Km) was determined by nonlinear regression to the Michaelis-Mention equation using GraphPad Prism (Version 10, GraphPad Software).

For compound testing in this assay, either 30 nM or 100 nM ABHD2^FL^ (depending on enzyme batch) was used, and samples were pre-incubated with 50 nl DMSO or 50 nl of 100-fold concentrated test compounds in DMSO for 15 min at room temperature prior to addition of RB substrate at a final concentration of 15 µM. Fluorescence measurement was performed after a 60 min incubation at room temperature.

#### ABHD2^L33–E425^

In initial compound screens, the effect of compounds on the enzymatic activity of ABHD2^L33-425^ was tested with *p*-nitrophenyl octanoate (*p*-NPO). For such assays, 50 nM ABHD2^L33-425^ was incubated with 500 µM *p*-NPO. Generally, 40 µl of protein solution was pre-incubated with a specific compound on ice for 30 min, transferred to a 96-well plate, and then placed in a plate reader with a thermostat function at 37 °C for 2 min. In parallel, buffer was incubated at 37 °C for 30 min, then *p*-NPO was added to the buffer, mixed, and 160 µl of substrate solution was added to each well containing 40 µl protein solution, thereby starting the reaction. This resulted in final assay conditions of 20 mM HEPES pH 8, 500 mM NaCl, 10% glycerol, 1 mM TCEP, 2% DMSO, 1.44% Methanol, and 500 µM *p*-NPO. Absorbance at 405 nm (of *p*-nitrophenol) was typically monitored for 10 min at 37 °C.

Enzymatic activity of ABHD2^L33-425^ was assessed using the fluorogenic substrate 7-HCA. Reactions (200 µl) were performed in 96-well plates with buffer containing 20 mM HEPES (pH 8.0), 500 mM NaCl, 10% (v/v) glycerol, 1 mM TCEP, and 2% (v/v) DMSO. For Michaelis–Menten kinetics, 5 nM ABHD2^L33-425^ was incubated for 20 min with varying concentrations of 7-HCA (µM): 0, 0.025, 0.05, 0.10, 0.25, 0.4, 0.5, 1, 2.5, 5, 7.5, and 10. Product formation was quantified against a standard curve of known umbelliferone concentrations by linear regression, and the Michaelis constant (Km) and V_max_ were determined by fitting the reaction rate against substrate concentrations via the Michaelis-Menten equation.

For compound testing, 40 µl of 50 nM ABHD2^L33-425^ was pre-incubated on ice for 30 min with 50 µM compound or DMSO in the same buffer, with all reactions normalized to DMSO. Plates were transferred to 37 °C for 5 min, and the enzymatic reaction was initiated by adding 160 µl of 37 °C pre-warmed substrate solution in identical buffer, to achieve final concentrations of 10 nM *ABHD2*^*L33–E425*^ and 10 µM 7-HCA. Initial screens were performed with compounds at 10 µM in duplicate. For IC50 determinations, compounds were spotted via echo-dispensing in triplicates at the following concentrations (µM): 0.00625, 0.00125, 0.0025, 0.0050, 0.0100, 0.0250, 0.500, 1, 5, 10, 50, and 100. Reactions were incubated for 10 min at 37 °C, and fluorescence was monitored every 30 s on a SpectraMax iD3 (Molecular Devices, LLC, San Jose, CA, USA), with excitation at 335 nm and emission at 450 nm.

For experiments with steroids and triterpenoids, final reactions (20 µl) were set up in 384-well plates with 10 nM ABHD2^L33–E425^, 5 µM steroid, and 300 nM 7-HCA in 20 mM HEPES (pH 8), 500 mM NaCl, 10 % (v/v) glycerol, 1 mM TCEP, and 0.4 % (v/v) DMSO. Preincubations were performed by mixing 40 µl of 50 nM ABHD2^L33–E425^ with 25 µM steroid triterpenoid, or DMSO on ice for 30 min in the same buffer, with samples normalized to the DMSO condition. Plates were then transferred to 37 °C for 10 min, and reactions were initiated by the addition of 4 µl of the enzyme-steroid, mixture to 16 µl of 37 °C pre-warmed substrate solution to achieve final concentrations of 10 nM ABHD2^L33–E425^, 5 µM steroid or triterpenoid, 300 nM 7-HCA, and 0.4 % (v/v) DMSO. Reactions were incubated for 10 min at 37 °C, and the fluorescence (excitation at 335 nm and emission at 450 nm) was monitored every 1 min on a BioTek Synergy H1 multimode reader (Agilent Technologies, Inc, Santa Clara, CA, USA). Percent activity was determined by comparing the rate of substrate production (slope) by ABHD2^L33–E425^ in the presence of steroids to the rate in the presence of DMSO, across three biological replicates. Statistical analysis was performed using GraphPad Prism 10.

#### ABHD10 and ABHD11

Enzymatic activities of ABHD10 and ABHD11 were measured in a colorimetric assay using p-nitrophenyl butyrate (*p*-NPB) as substrate. Enzymes at 1500 nM ABHD10 and 125 nM ABHD11 in 20 mM HEPES (pH 7.5), 150 mM NaCl, and 2% (v/v) DMSO were pre-incubated in 96-well plates with 50 µM test compound for 30 min at 4°C. Plates were then equilibrated at 37°C for 2 min, and reactions were initiated by adding 160 µL of pre-warmed substrate solution (same buffer with 625 µM *p*-NPB) to a total volume of 200 µl, yielding final assay concentrations of 300 nM ABHD10, 25 nM ABHD11, and 500 µM pNPB. The substrate solution was prepared immediately before addition to the enzyme mixtures. Absorbance at 405 nm was recorded on a SpectraMax iD3 plate reader (Molecular Devices) for 20 min with continuous monitoring at 30 s intervals.

### Analysis of steroid and compound binding to purified ABHD2

We used differential scanning fluorimetry (DSF) to investigate the interaction of steroids and compounds with recombinant ABHD2. To this end, 4 µM of recombinant ABHD2 proteins were incubated on ice for 30 min with steroids or test compounds at 20 µM or 40 µM in the respective final purification buffer. Vehicle controls were 5 % (v/v) DMSO for ABHD2^FL^ experiments and 2 % or 4 % (v/v) DMSO for ABHD2^L33–E425^. Following incubation, samples were spun at 15,000 × g and loaded into Prometheus− NT.48 capillaries (NanoTemper Technologies GmbH, Munich, Germany). After a 5 min equilibration at 25 °C, samples were heated to 95 °C with a linear ramp of 1 °C/min on a Prometheus NT.48 instrument. Fluorescence emission at 330 and 350 nm was recorded in triplicate, and protein melting temperatures were determined as inflection points of a sigmoidal fit to the 350/330 nm fluorescence ratio as a function of temperature. An increase in the protein melting temperatures, i.e., enhanced resistance against heat-induced denaturation, is indicative of ligand/compound binding to the protein.

### Analysis of compound binding to ABHD2 in intact cells

We used Cellular Thermal Shift Assays (CETSA) to study the binding of compounds to ABHD2 in intact cells. To this end, HEK293T cells were seeded in 6-well plates at 1 × 10^6^ per well, and reverse-transfected with 0.2 µg of N-terminally HiBIT–tagged ABHD2^L33-425^ and 1.8 µg of empty plasmid using X-tremeGene XP transfection reagent, following the manufacturer’s instructions. The following day, cells were trypsinised and resuspended in OptiMEM (without phenol red) at a density of 2 × 10^5^ cells/ml. After addition of test compounds or DMSO, cells were transferred into 96-well PCR plates (40 µl per well), incubated for 1 h at 37 °C, and heated in a thermocycler for 3 min at temperatures between 37°C and 60°C. After an additional 3 min at room temperature, 40 µl of LgBIT solution (200 nM LgBIT, 2 % (v/v) NP-40, and protease inhibitors in OptiMEM) was added and incubated for 10 min at room temperature. Subsequently, 20 µl of NanoGlo substrate (8 µl/ml, Promega Corporation, Madison, WI, USA) was added, mixed gently, and 20 µl was transferred to 384-well white plates. Bioluminescence was read with a ClarioStar plus plate reader (BMG LABTECH). Recovery of soluble protein after heating was assessed by luminescence detection of reconstituted luciferase, with signals normalised to non-heated controls kept at 37 °C. For multiple compound profiling, a single heating temperature of 50 °C was used, and normalised to cells incubated at 37 °C. An increase in soluble protein after heating, i.e., enhanced resistance against heat-induced denaturation, is indicative of ligand/compound binding to the protein.

### Sperm intracellular Ca^2+^ measurements

Measurements of the changes in sperm intracellular Ca^2+^ concentration ([Ca^2+^]_i_) were performed on motile human sperm purified by swim-up. Semen samples were allowed to liquefy for 15–60 min at 37 °C, and 0.5–1 ml liquified semen was layered beneath 4 ml human tubal fluid medium (HTF; 97.8 mM NaCl, 4.69 mM KCl, 0.2 mM MgSO_4_, 0.37 mM KH_2_PO_4_, 2.04 mM CaCl_2_, 2.78 mM glucose, 0.33 mM sodium pyruvate, 21.4 mM lactic acid, 4 mM NaHCO_3_, and 21 mM HEPES, pH 7.35 adjusted with NaOH) in 50 ml conical tubes. Motile sperm were allowed to swim up into the HTF for 60–90 min at 37°C, collected, washed twice (700 × g, 20 min, 37°C), and resuspended in HTF containing 3 mg/ml HSA at a density of 1 × 10^7^ sperm/ml. Changes of [Ca^2+^]_i_ in motile human sperm were measured as previously described (Rennhack et al., 2018), using a fluorescence plate-reader assay (FLUOstar Omega, BMG LABTECH) with 384-well plates. Swim-up-purified sperm were loaded with Fluo4-AM (5 μM, Thermo Fisher Scientific, Waltham, MA, USA) and Pluronic F-127 (0.05 % w/v) for 20 min at 37 °C. Excess dye was removed by centrifugation at 700 × g for 5 min, supernatant removal, and resuspension to 5 × 10^6^ sperm/ml in HTF.

For recordings, wells of a 384-well plate were filled with 50 μl of the dye-loaded sperm suspension. Fluorescence was excited at 480 nm, and emission recorded at 520 nm using a plate reader (Fluostar Omega, BMG Labtech). After establishing a baseline for at least 60 s, 25 µl test compound solutions were added and the fluorescence response recorded. After 4 min, 7.5 µl progesterone was added to reach a final concentration of 3 µM and ensuing changes in fluorescence were recorded. For analysis of compound-evoked Ca^2+^ signal, basal fluorescence (F_0_) was determined prior to compound addition. For determination of inhibitory effects of compounds on progesterone-evoked Ca^2+^ signals, basal fluorescence (F_0_) was determined before progesterone addition. The CatSper inhibitor TS150 (Schierling et al, 2022) was used as control. Each compound was tested in technical duplicates across ≥3 biological replicates, using sperm from at least two donors and performed on at least two separate days.

### Sperm hyperactivation assay

Sperm hyperactivation was assessed in human sperm purified by density gradient centrifugation. Semen samples (n = 7) were allowed to liquefy for 15–30 min at 37 °C, and up to 2 ml liquified semen was layered onto a two-step GM501 gradient system (45 %/90 %; Gynemed; Sierksdorf, Germany) pre-equilibrated at 37 °C. Gradients were centrifuged for 20 min at 400 × g (no break). The 90% pellet was collected, washed in SpermActive medium (Gynemed; composition per manufacturer: NaCl, KCl, MgSO_4_.7H_2_O, KH_2_PO4, 1.77 mM CaCl_2_.2H_2_O, D(+)-glucose anhydrous, sodium pyruvate, sodium lactate, EDTA, alanyl-glutamine, essential and non-essential amino acids, 25.71 mM NaHCO_3_, 15 mM HEPES, 5 g/l human serum albumin, 10 mg/l gentamicin and phenolred), and incubated for at least 2 h at 37 °C, 5 % CO_2_.

For compound testing, sperm suspensions with 7.5 x 10^6^ sperm/ml in SpermActive were treated with either DMSO (vehicle) or 10 µM test compound. For basal and induced hyperactivation conditions, DMSO or progesterone (3 µM), respectively, was added. Samples were incubated 30 min at 37 °C, 5 % CO_2,_ before motility analysis.

Motility was analysed using a CASA system (CEROSII− Animal Motility, software version 1.16; Hamilton Thorne, Beverly, MA, USA), mounted on an Axiolab 5 microscope (Zeiss, Germany), equipped with a TPi-SX Thermal Plate (Tokai Hit; Shizuoka-ken, Japan) and a JAI CM-040-GE CCD camera (JAI; Tokio, Japan). Leja slides (20 µm chamber depth; Leja Products B.V., Nieuw-Vennep, The Netherlands) were pre-warmed to 37°C, cleaned with lens paper, and loaded by capillary action with 6 µl sperm suspension; capillary correction was set to 1.3. Sperm tracks were recorded at 37°C using a 10× negative-phase objective, acquiring 60 frames at 60 Hz with 4 ms exposure and gain set to 600. For each sample and condition, 300–500 motile sperm from at least 10 non-overlapping fields were analysed. Hyperactive sperm were identified using the SORT function with the following Boolean argument: VCL > 150 µm/s and LIN < 50% and ALH > 7 µm. For each sample and condition, the percentage of hyperactive sperm was normalised to the percentage of motile sperm, and the mean of two technical duplicates was reported.

### Statistical analysis

#### ABHD2^L33-E425^ enzymatic activity

Statistical analysis was performed using GraphPad Prism 10 for MacOS (version 10.3.1). IC_50_ values were determined by a four-parameter dose response curve using a variable Hill slope.

#### Ca^2+^ assay

Data analysis and graphical representation were performed with GraphPad Prism software version 10.6.1 (San Diego, CA, USA). Changes in Fluo-4 fluorescence are depicted as ΔF/F_0_ (%), i.e. the change in fluorescence (ΔF) relative to the mean basal fluorescence (F_0_) before application of buffer or compounds/ligands, to correct for intra- and inter-experimental variations in basal fluorescence among individual wells or to correct for compound-evoked changes in fluorescence. Signal amplitudes evoked in the presence of compounds were normalized to the maximal signal amplitude evoked by 3 µM progesterone in the absence of the compound (set to 1) to ease comparisons between experiments.

#### Hyperactivation assay

Statistical analysis was performed using GraphPad Prism 10 for Windows (version 10.6.1). Data normality was assessed using Shapiro-Wilk test. Differences between basal and induced hyperactivation were evaluated using a paired t-test, with effective pairing confirmed across all comparisons. *P* < 0.05 was considered statistically significant.

## Results

### Compound 1 inhibits ABHD2 activity

To evaluate the inhibitory effects of small molecules on human ABHD2, we expressed full-length human ABHD2 in SF9 cells and purified the protein (ABHD2^FL^; Fig. 1, Supplementary Table S1). The enzymatic activity of ABHD2^FL^ was quantified using two complementary fluorogenic substrates: 7-hydroxycoumarinyl arachidonate (7-HCA; Supplementary Figure S1A) and resorufin butyrate (RB; Supplementary Figure S1B). Both 7-HCA and RB are established substrates for measuring monoacylglycerol lipase (MAGL) activity (Miceli et al., 2019, Wang et al., 2008). Given the predicted functional similarity between MAGL and ABHD2, we hypothesized that ABHD2^FL^ would also hydrolyse these substrates. Consistent with our hypothesis, ABHD2^FL^ catalysed the hydrolysis of both 7-HCA and RB, with K_m_ values of 59 µM and 23 µM, respectively (Fig. 2A, B; Supplementary Figure S1C, D). To validate the assay, we profiled known, unselective lipase inhibitors. Both MAFP (Amara et al., 2012) and Orlistat (Navia-Paldanius et al., 2012) (Parkkari et al., 2014) inhibited ABHD2^FL^ activity in a dose-dependent manner (Fig. 2 C), with IC_50_ values of 51 nM and 969 nM, respectively, in the 7-HCA assay, and 44 nM and 305 nM, respectively, in the RB assay. Baggelar *et al*. (2019) reported that Compound **1** inhibits ABHD2 in lysates of cells overexpressing the protein. Using the recombinant protein, we confirmed that Compound **1** inhibits ABHD2^FL^ with IC_50_ values of 316 nM (pIC_50_ = 6.50 ± 0.14, n = 4) in the 7-HCA assay and 136 nM (pIC_50_ = 6.87 ± 0.14, n = 3) in the RB assay (Fig. 2D).

**Figure 1.**
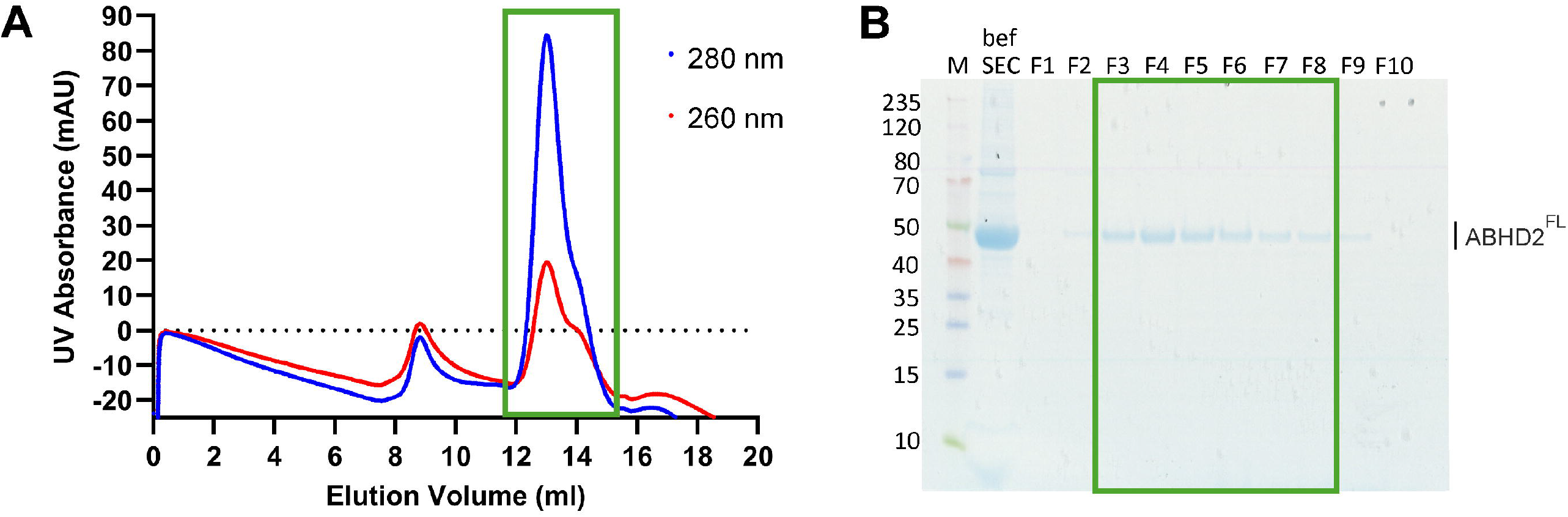
Full-length ABHD2 (ABHD2^FL^) purification. **A)** Size-exclusion chromatography (SEC) trace from the final purification step of recombinant ABHD2^FL^. Absorptions at 280 nm (blue) and 260 nm (red) are shown in arbitrary milli-absorbance units (mAU) against elution volume. The main peak selected for pooling and concentration is indicated by the green box. **B)** SDS-PAGE analysis of the final SEC purification, showing the protein before gel filtration (bef SEC) and the indicated fractions (F1-F10) of the gel filtration run. Fractions highlighted in the green box were pooled, concentrated and frozen to generate the final protein batch.

**Figure 2.**
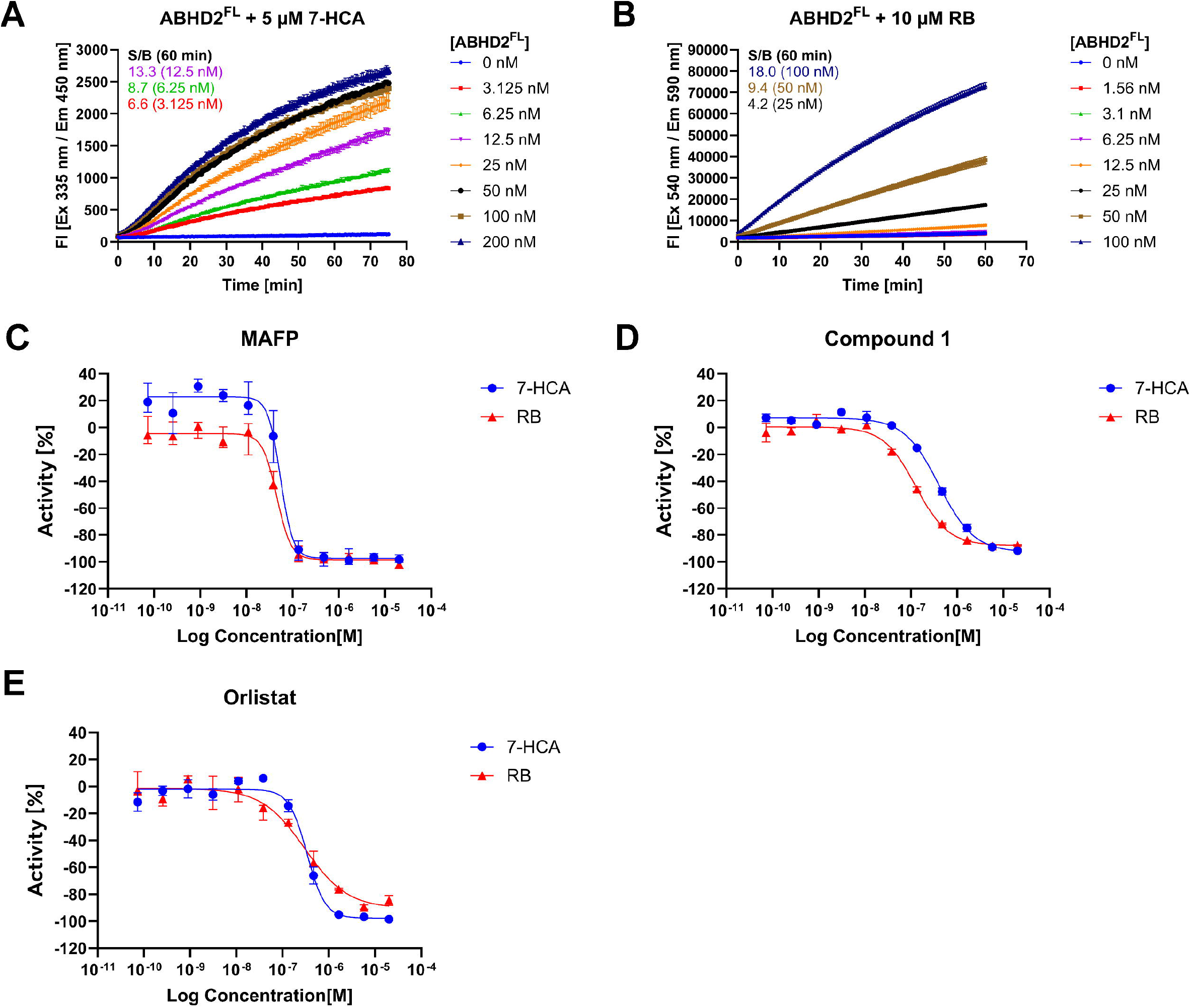
Enzymatic activity of purified full length ABHD2 (ABHD2^FL^) can be inhibited by lipase inhibitors and Compound 1. **A)** The indicated ABHD2^FL^ concentrations were incubated with 5 µM 7-hydroxycoumarinyl arachidonate (7-HCA) substrate in assay buffer for 75 min at RT. Fluorescence was monitored every 30 s at excitation and emission wavelengths of 335 and 450 nm, respectively. Signal-to-background (S/B) ratios were calculated for 3.1 (red), 6.25 (green) and 12.5 nM (purple) ABHD2^FL^. **B)** The indicated ABHD2^FL^ concentrations were incubated with 10 µM resorufin butyrate (RB) substrate in assay buffer for 60 min at RT. Fluorescence was monitored every 30 s at excitation and emission wavelengths of 540 and 590 nm, respectively. Signal-to-background (S/B) ratios were calculated for 25 (black), 50 (brown) and 100 nM (blue) ABHD2^FL^. **C), D)** Dose-response curves were generated by incubating a fixed ABHD2^FL^ concentration with 5 µM 7-HCA (blue curve) or 15 µM RB (red curve) substrate in the presence of increasing concentrations of methyl arachidonyl fluorophosphonate (MAFP), Orlistat and Compound 1. Percent activity was plotted against log10 inhibitor concentration and fitted by nonlinear regression to obtain IC_50_ values. For each measurement, fluorescence intensity values were normalized to neutral controls minus inhibitors using a Standard Two-Point Normalization (Genedata Screener), with the neutral control (DMSO only, full activity) set to 0% and the inhibitor control (no enzyme, no activity) set to -100%.

### Compound 1 binds ABHD2

In addition to enzyme-activity assays, we also wanted to study the binding of ligands and compounds to recombinant ABHD2 through Differential Scanning Fluorimetry (DSF) and to ABHD2 in intact cells through Cellular Thermal Shift Assays (CETSA). Both techniques infer binding from ligand/compound-induced changes in thermal protein stability. Yet, structural predictions and sequence analysis indicate that the N-terminal residues 10–30 of ABHD2 form an alpha helix consistent with a transmembrane anchor that may confound thermal ABHD2 denaturation as a readout in intact cells. We therefore also expressed and purified a truncated variant lacking the first 32 N-terminal residues produced in *E. coli*, which we refer to as ABHD2^L33–E425^ (Supplementary Figure S2). We investigated whether the enzymatic activity and the action of known inhibitors is preserved in ABHD2^L33–E42^ The truncated protein indeed retained enzymatic activity with 7-HCA as substrate, exhibiting a Km of 280 nM (Supplementary Figure S3 A, B). MAFP inhibited ABHD2^L33-E425^ with IC_50_ values of 4.6 nM, and Compound **1** with an IC_50_ of 1.3 µM (Supplementary Figure S3C). This indicates that not only full-length ABHD2^FL^ but also truncated ABHD2^L33–E425^ is suitable for studying ligand/compound binding to the recombinant, purified protein through DSF, and, moreover, that ABHD2^L33–E425^ can be used instead of ABHD2^FL^ to study ligand/compound binding in intact cells through CETSA.

### Steroids and triterpenoids neither bind to ABHD2 nor modulate its enzymatic activity

To determine whether steroids or structurally related triterpenoids affect ABHD2 activity, we studied the action of progesterone, cholesterol, testosterone, pregnenolone sulfate, hydrocortisone, 17β-estradiol, and the plant-derived triterpenoids pristimerin and lupeol on its enzymatic activity. Except for pregnenolone sulfate, the steroids did not alter the enzymatic activity of purified ABHD2^FL^ or ABHD2^L33–E425^ (Supplementary Figure S4). Yet, pregnenolone sulfate exhibited only a weak inhibitory action on ABHD2^FL^, but not on ABHD2^L33–E425^, at concentrations ≥ 5 µM. Pristimerin did not affect ABHD2 activity, whereas at ≥ 5 µM, lupeol exhibited a weak inhibitory action on ABHD2^L33–E425^ (Supplementary Figure S4 C) but not on ABHD2^FL^ (Supplementary Figure S4 A, B). The weak inhibitory actions of pregnenolone sulfate and lupeol at high micromolar concentrations might reflect non-specific effects of the compounds in the assay rather than a genuine inhibitory action on ABHD2.

To scrutinize the interaction of the steroids and the triterpenoids with ABHD2, their potential binding to purified ABHD2 was assessed by DSF. Pre-incubation with progesterone, cholesterol, testosterone, pregnenolone sulfate, hydrocortisone, or lupeol did not affect heat-induced denaturation of ABHD2^L33–E425^ (Supplementary Figure S5A). DSF analysis for 17β-estradiol was inconclusive due to fluorescence interference. Consistently, progesterone did not affect heat-induced denaturation of ABHD2^FL^ either, whereas Compound **1**, included as a positive control, increased the melting temperature by 6 °C (Supplementary Figure S5B). Together, these findings indicate that steroids and steroid-like triterpenoids do not bind to ABHD2, explaining their failure to suppress its enzymatic activity.

### Design, synthesis, and characterization of novel selective ABHD2 inhibitors

Having confirmed that Compound **1** inhibits both full-length and truncated ABHD2, we next sought to enhance its potency and selectivity through structure-based optimization. We designed and either purchased or synthesized 64 derivatives of Compound **1** and screened them for binding to purified ABHD2^L33–E425^ and to ABHD2^L33–E425^ in HEK293T cells (Fig. 3A, B); we also assessed the inhibition of enzymatic activity of purified ABHD2^L33–E425^ and ABHD2^FL^ (Fig. 3A, Supplementary Figure S6A-C, Supplementary Table 2).

**Figure 3.**
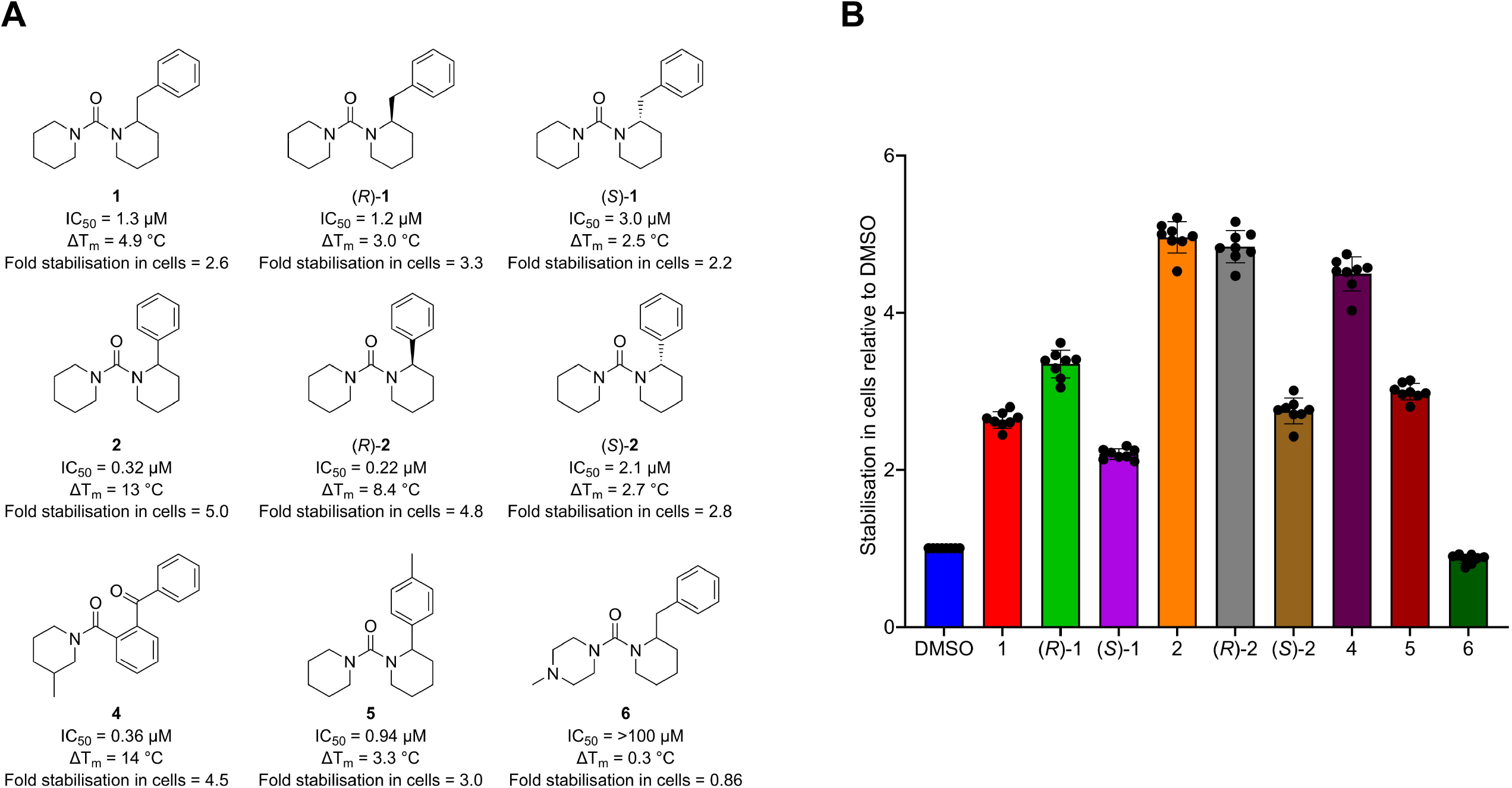
Characterisation of ABHD2 inhibitors. **A)** Chemical structures of compounds **1, (*R*)-1, (*S*)-1, 2, (*R*)-2, (*S*)-2, 4, 5**, and **6** annotated with IC_50_ values for ABHD2^L33-425^ assessed by a fluorometric assay using 7-HCA as substrate, ΔT_m_ shifts for ABHD2^L33-425^ determined by DSF, and fold cellular stabilisation of ABHD2^L33-425^ in HEK293T cells. **B)** Cellular target-ligand interaction in HEK293T cells with ABHD2^L33-425^ measured as a recovery of signal when heating from 37 °C to 50 °C, relative to DMSO (i.e. DMSO = 1.0).

Compounds **2, 4** and **5**, which showed similar to modestly improved potency to inhibit enzymatic activity of ABHD2^FL^ and ABHD2^33–E425^ compared to Compound **1** (Fig. 3A; Supplementary Figure S6A-C), were selected for further evaluation. A structurally related but inactive compound (Compound **6**) served as negative control. Moreover, Compounds **1** and **2**, initially tested as racemic mixtures, were resolved into enantiomers. The (*R*)-enantiomers displayed higher potency than the (*S*)-enantiomers (Fig. 3A; Supplementary Table 2; Supplementary Figure S6B, C), consistent with stereospecific binding to a chiral binding pocket.

In the DSF assay, compounds **2, 4** and **5** also increased the thermal stability of purified ABHD2^L33-E425^ by 3 - 14 °C, confirming that the compounds bind to ABHD2, whereas the negative control (Compound **6**) had no effect (Fig. 3A, Supplementary Figure S7). Moreover, compounds 2 and 4 also exhibited high selectivity, showing no detectable inhibition of the related ABHD family members ABHD10 and ABHD11 (Supplementary Figure S8). We did not test the action of compound 5 on ABHD10 or ABHD11.

To evaluate the binding of the compounds to ABHD2 in intact cells, we performed CETSA using HEK293T cells transfected with N-terminally HiBiT-tagged ABHD2^L33-E425^. Cells were treated with the respective compound or the vehicle (DMSO), heated to 50 °C, and the fraction of soluble ABHD2^L33-E425^ relative to that observed at 37 °C (reference) was determined as a measure of heat-induced denaturation. All active compounds, including **1**, (*R*)**-1**, (*S*)**-1**, 2, (*R*)**-2**, (*S*)**-2, 4** and **5**, reduced the heat-induced denaturation of ABHD2, i.e., the compounds increased the fraction of soluble protein at 50 °C, with fold-changes ranging from 2.2 to 5.0 relative to the vehicle control (Fig. 3B). The most pronounced action was observed for compounds **2**, (*R*)**-2**, and **4**, consistent with their in *vitro* potency and suggesting efficient cell penetration and ABHD2 binding.

### ABHD2 inhibition does not affect progesterone-induced Ca^2+^ influx or hyperactivation in human sperm

Finally, to test whether ABHD2 activity is required for progesterone-induced Ca^2+^ influx through CatSper and resulting sperm hyperactivation, we evaluated the effects of ABHD2 inhibitors in human sperm.

Changes in the intracellular Ca^2+^ concentration ([Ca^2+^]_i_) of human sperm were monitored using the fluorescent Ca^2+^ indicator Fluo-4. Challenging sperm with the ABHD2 inhibitors *(R)*-**1**, (*R*)-**2, 4**, or the inactive control Compound **6**, caused only miniscule changes in [Ca^2+^]_i._ Moreover, the amplitude and waveform of progesterone-induced Ca^2+^ signals were similar in the presence of the ABHD2 inhibitors and Compound **6** (Fig. 4 A). In contrast, the CatSper inhibitor TS150 (Schierling et al., 2023) completely inhibited the progesterone-induced Ca^2+^ signal, confirming assay specificity.

**Figure 4.**
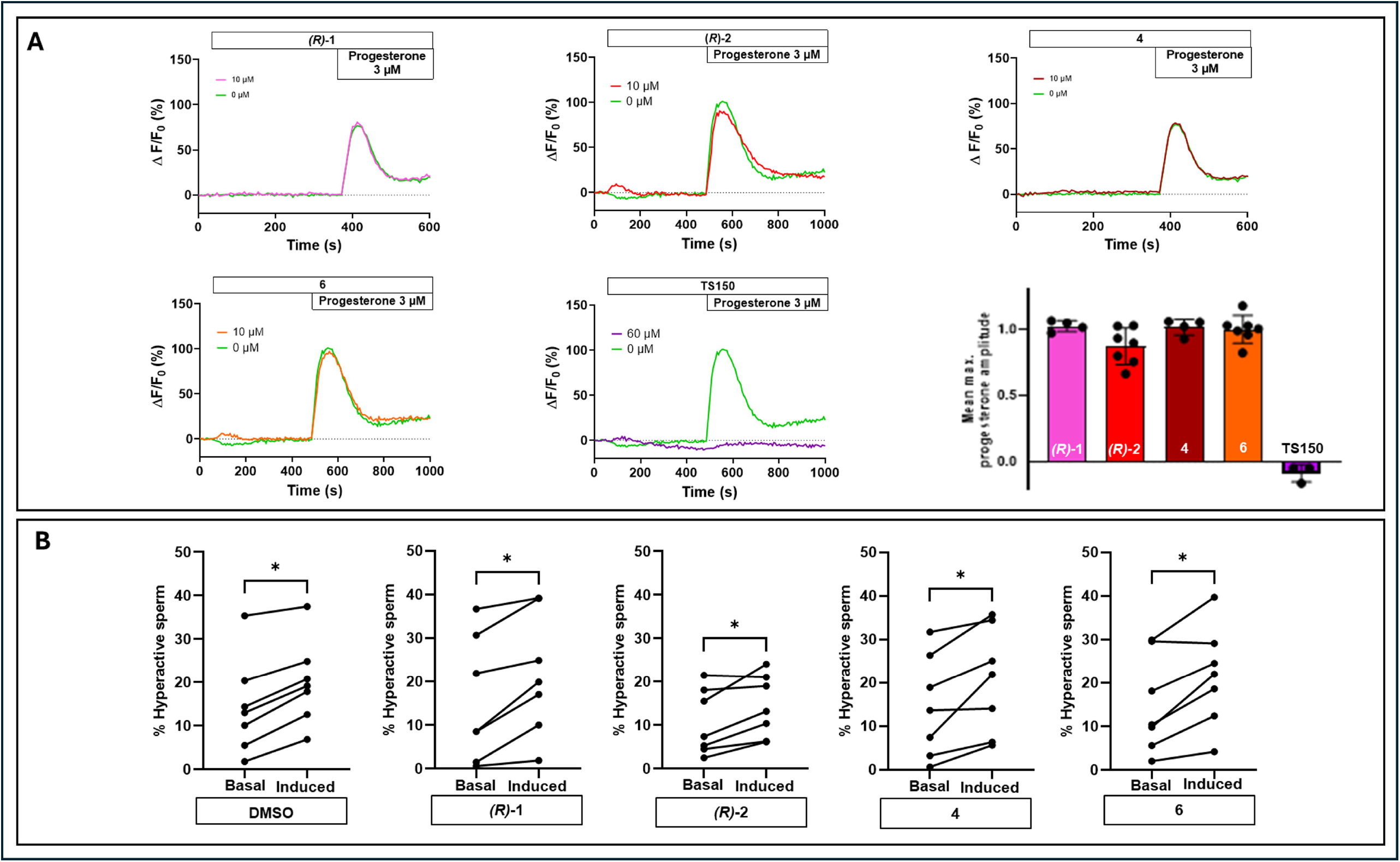
ABHD2 inhibitors have no effect on progesterone-evoked Ca^2+^ signals or hyperactivation in human sperm. **A)** Representative Ca^2+^ signals in human sperm loaded with a fluorescent Ca^2+^ indicator evoked by the application of buffer or compounds **(*R*)-1, (*R*)-2, 4** and **6** (10 µM), or TS150 (60 µM) and subsequent stimulation by progesterone (3 µM). ΔF/F (%) indicates the percentage change in fluorescence (ΔF) with respect to the mean basal fluorescence (F) before application of the respective compound. The bar graph (right) shows mean (± SD) maximal signal amplitude evoked by progesterone (3 µM) in sperm pre-incubated with compound relative to that evoked in its absence (set to 1). **B)** Paired plots comparing basal and progesterone-induced sperm hyperactivation. Sperm suspensions were treated with either DMSO (vehicle) or 10 µM test compound (**(*R*)-1, (*R*)-2, 4** and **6**). Induced conditions (3 µM progesterone) were compared to basal conditions (DMSO). For each of 7 independent experiments, the mean percentage of hyperactive sperm (normalized to motile sperm) across two technical replicates is shown. Statistically significant differences between basal and induced conditions are indicated by an asterisk (*, P < 0.05).

Hyperactivation of human sperm was assessed using computer-assisted sperm analysis (CASA). Compounds *(R)*-**1** (*R*)-**2, 4**, and **6** did not affect basal hyperactivation (Supplementary Figure S9). Stimulation of sperm with progesterone increased the fraction of hyperactivated sperm. In line with the results from [Ca^2+^]_i_ recordings, progesterone-induced hyperactivation was not affected by the ABHD2 inhibitors or Compound **6** (Fig. 4 B).

Taken together, these findings demonstrate that ABHD2 activity is not required for progesterone-evoked Ca^2+^ influx through CatSper and resulting hyperactivation, refuting the model of non-genomic progesterone action in human sperm proposed by Miller and colleagues (Miller, et al., 2016).

## Discussion

Our study demonstrates that ABHD2 activity is dispensable for the progesterone-driven events in human sperm, Ca^2+^ influx and hyperactivation. Potent, selective, and cell-active ABHD2 inhibitors do not affect the Ca^2+^ and motility responses evoked by progesterone. In line with that finding, we show that steroids and steroid-like triterpenoids neither bind to nor modulate the enzymatic activity of ABHD2. Together, these results refute the prevailing hypothesis that ABHD2 mediates non-genomic steroid signalling in sperm and indicate that a different molecular mechanism underlies CatSper activation by steroids.

A key strength of our study is the rigorous biochemical characterisation of ABHD2. Prior studies have reported recombinant expression of ABHD2 in heterologous systems (Baggelaar, et al., 2019, M et al., 2016, Miller, et al., 2016), yet data on protein purity and a rigorous quantitative characterisation of enzymatic activity were lacking. Here, we isolated two recombinant ABHD2 proteins to high purity and established powerful, quantitative biochemical assays for monitoring their activity. These advances enabled us to directly test whether steroids and/or triterpenoids bind ABHD2 or modulate its activity, and to determine whether small molecules can inhibit ABHD2 in cells.

The mechanism driving CatSper activation and the subsequent Ca^2+^ flux essential for sperm function remains elusive. CatSper is not only activated by progesterone but also by various other steroids that are present in female reproductive fluids (Brenker, et al., 2018, De Toni et al., 2023, Jeschke et al., 2021, Lishko, et al., 2011, Mannowetz, et al., 2017, Rehfeld, 2020, Strunker, et al., 2011). The current model of the non-genomic activation of CatSper by steroids suggests that hormones bind to ABHD2, triggering hydrolysis of 2-arachidonoylglycerol (2-AG), thereby relieving CatSper inhibition and facilitating Ca^2+^ influx and sperm hyperactivation (Miller, et al., 2016). Our findings do not support this mechanism. Across different assay formats, ABHD2 enzymatic activity was unaffected not only by progesterone but also other steroids known to activate CatSper, i.e, pregnenolone sulfate, testosterone, hydrocortisone, and cholesterol (Brenker, et al, 2018, Rehfeld 2020, Jeschke, et al. 2021). Notably, the action of the steroids was evaluated at concentrations spanning and exceeding physiological ranges. Furthermore, we did not detect binding of steroids to purified ABHD2, explaining the failure of the steroids to interfere with enzyme activity. Moreover, reports on the effects of plant-derived triterpenoids such as pristimerin and lupeol on CatSper activation are conflicting (compare Brenker et al., 2018 and Rehfeld et al., 2020 with Mannowetz, et al, 2017). We show that pristimerin and lupeol do not suppress ABHD2 activity and/or bind to purified t ABHD2. Collectively, these data indicate that ABHD2 activity is not modulated by steroids or triterpenoids and that steroid-induced CatSper activation occurs through a mechanism independent of ABHD2.

The previously suggested role of ABHD2 in CatSper regulation was inferred from experiments using the irreversible, broad-spectrum serine hydrolase inhibitor MAFP (Miller, et al., 2016). In that study, MAFP suppressed progesterone-induced Ca^2+^ influx and motility changes. However, MAFP lacks specificity for ABHD2, raising the possibility that these effects were mediated via other serine hydrolases and/or off-target effects of the compound. Using potent and selective ABHD2 inhibitors, we demonstrate that inhibition of ABHD2 activity does neither impair progesterone-induced Ca^2+^ influx through CatSper nor sperm hyperactivation, thereby confirming that ABHD2 activity is dispensable for the non-genomic action of steroids in human sperm.

In summary, our findings refute the model that ABHD2 serves as a non-genomic progesterone receptor. Although the molecular basis of steroid-induced CatSper activation remains unresolved, our data demonstrate unequivocally that this process does not involve ABHD2. Consequently, ABHD2 does not play a critical role in sperm hyperactivation and is unlikely to represent a viable target for the development of non-hormonal contraceptives. Identifying the true molecular mediators of CatSper activation will be essential for advancing both fundamental reproductive biology and contraceptive research.

## Data availability statement

The data underlying this article are available in the article and in its online supplementary material.

## Authors’ Role Statement

M.E., E.M.C., and O.A. designed and executed experiments with ABHD2^L33-E425^, including compound and steroid testing, IC_50_ determination, analysed the resulting data and contributed to manuscript writing. A.A. designed and performed the hyperactivation assays, analysed the resulting data and was responsible for drafting parts of the manuscript. K.V. designed and performed the expression and purification of ABHD2^FL^ and the ABHD2^FL^ nanoDSF measurement and contributed to manuscript writing. A.T. designed and supervised all experiments involving recombinant ABHD2^FL^, including enzymatic activity assays and IC_50_ determinations of compounds and steroids, analysed the associated data and contributed to manuscript writing. N.D., A.H., and M.W. conducted all experiments involving recombinant ABHD2^FL^, including enzymatic activity assays and IC_50_ determinations of compounds and steroids. N.T.-E. and E.W. expressed and purified recombinant ABHD2^L33-E425^. P.R. and K.B. designed and synthesized the derivatives, with oversight of compound synthesis by E.H. and M.H., P.L., M.S., M.A.C., and D.B.-L. designed and performed the cellular thermal shift assays. W.F.Z. and J.M. designed and performed the functional Ca^2+^ influx assays and analysed the data and contributed to manuscript writing. L.T., C.B., and T.S. supervised experimental work and contributed to data interpretation. M.H.T., M.S., and R.L. contributed to study planning and supervision of specific experimental components and manuscript writing. A.M.E., O.G., RL, and C.T. conceptualized the study. C.T. planned, supervised and coordinated the overall project and led manuscript writing.

All authors contributed to interpretation of the findings, critically revised the manuscript for important intellectual content, approved the final version to be published, and agree to be accountable for all aspects of the work.

