## Supplemental Figures S1-S9 for "ABHD2 activity is not required for the non-genomic action of progesterone on human sperm"

### Supplementary figures

Madison Edwards[1*], Alexandra Amaral[2*], Eve M. Carter[3*], Oliver Arnolds[4*], Karen Vester[2], Anna Thrun[2], Edvard Wigren[4], Evert Homan[5], Pauline Ribera[6], Kirsty Bentley[5], Martin Haraldsson[6], Nmesoma Theo-Emegano[1], Peter Loppnau[1], Magdalena M. Szewczyk[1], Michelle A. Cao[1], Dalia Barsyte-Lovejoy[1], Nicole Dittmar[2], Anika Hans[2], Mandy Weber[2], Jens Münchow[7,8], W. Felix Zhu[7,8], Louisa Temme[8,9], Christoph Brenker[10], Timo Strünker[8,10], Michael Sundström[4], Matthew H. Todd[3], Aled M. Edwards[1], Ralf Lesche[2], Opher Gileadi[4¶], Claudia Tredup[11,12¶]

**Supplementary Figure S1. Full length ABHD2 (ABHD2$^{FL}$) hydrolyzes the fluorogenic substrates 7-hydroxycoumarinyl arachidonate (7-HCA) and resorufin butyrate (RB). A)** Molecular structure of 7-hydroxycoumarinyl arachidonate (7-HCA). **B)** Molecular structure of resorufin butyrate (RB). **C)** The indicated 7-HCA concentrations were incubated with 6 nM ABHD2$^{FL}$ in assay buffer for 60 min at RT. Fluorescence was monitored every 30 sec at excitation and emission wavelengths of 335 and 450 nm, respectively. Michaelis constant (Km) was determined by nonlinear regression to the Michaelis-Mention equation. **D)** The indicated RB concentrations were incubated with 25 nM ABHD2$^{FL}$ in assay buffer for 60 min at RT. Fluorescence was monitored every 30 sec at excitation and emission wavelengths of 540 and 590 nm, respectively. Michaelis constant ($K_m$) was determined by nonlinear regression to the Michaelis-Mention equation.

**Supplementary Figure S2. Truncated ABHD2 (ABHD2$^{L33\text{-}E425}$) purification. A)** Domain structure of human ABHD2. For the recombinant protein, the putative transmembrane (TM) helix (amino acids 1-32) was removed. **B)** SDS-PAGE analysis of purified ABHD2$^{L33\text{-}E425}$ after Size Exclusion Chromatography (SEC) shows the 45 kDa protein. **C)** SEC trace of absorbance at 280 nm (mAU) vs. elution volume (mL) from the final purification step of recombinant ABHD2$^{L33\text{-}E425}$ (blue). The SEC trace for ABHD2$^{L33\text{-}E425}$ is overlayed with the SEC trace for BioRad Chromotography Standards (orange), which were run on the same column in a separate experiment. Comparing the elution volume of ABHD2$^{L33\text{-}E425}$ to the standards allows for a predicted molecular weight which most closely corresponds to a tetramer.

**Supplementary Figure S3. Hydrolase activity of ABHD2$^{L33\text{-}E425}$. A)** The hydrolase activity of ABHD2$^{L33\text{-}E425}$ was assessed by a fluorometric assay using 7-hydroxycoumarinyl arachidonate (7-HCA) as substrate. A $K_m$ and $k_{cat}$ of 280 nM and 2.8 pmol substrate/min/pmol enzyme, respectively, were calculated, from **B)** the original Km curves. **C)** ABHD2$^{L33\text{-}E425}$ is inhibited by MAFP with an IC50 of 4.6 nM.

**Supplementary Figure S4. Steroid hormones and triterpenoids do not modulate the enzymatic activity of ABHD2. A), B)** Dose-response curves were generated by incubating a fixed ABHD2$^{FL}$ concentration with 5 μM 7-HCA (**A**) or 15 μM RB (**B**) substrate in the presence of increasing concentrations of positive controls Compound 1 and Orlistat, the indicated steroid hormones or triterpenoids. Percent activity was plotted against log10 inhibitor concentration. **C**) Percent activity of 10 nM ABHD2$^{L33–E425}$ in the presence of 5 μM steroids, triterpenoids, or MAFP compared to DMSO control. Percent activity is a comparison of the rate of substrate production by ABHD2$^{L33–E425}$ in the presence of steroids or triterpenoids to the rate in the presence of DMSO, across three biological replicates.

**Supplementary Figure S5. Binding of steroids to ABHD2. A)** Binding of steroids to ABHD2$^{L33-E425}$ was assessed using differential scanning fluorimetry (DSF) following 30 min pre-incubation with 40 μM of the indicated steroids. Dashed lines denote the middle of each melting curve in the fluorescence plot, which corresponds to the peak of the first derivative curve. These lines mark the melting temperature of each sample on the X-axis. All samples showed the same melting temperature, indicating that no binding was observed in the presence of progesterone, cholesterol, testosterone, pregnenolone sulfate, hydrocortisone, lupeol, or 17α-hydroxyprogesterone. **B)** ABHD2$^{FL}$ melting curve assessment in the presence of 20 μM progesterone or compound **1**.

**Supplementary Figure S6. Characterisation of ABHD2 inhibitors. A)** Enzymatic activity of ABHD2$^{FL}$ and ABHD2$^{L33-E425}$ was assessed in the presence of Compound **1** derivatives. Percent inhibition of ABHD2$^{FL}$ in the 7-HCA assay at 20 μM compound concentration was correlated with percent inhibition of ABHD2$^{L33-E425}$ in pNP octanoate assay at 10 μM compound concentration. **B), C)** Dose-response curves were generated by incubating a fixed ABHD2$^{FL}$ (**B**) or ABHD2$^{L33-E425}$ (**C**) concentration with 7-HCA substrate in the presence of increasing concentrations of the indicated compounds. Percent activity was plotted against log10 inhibitor concentration and fitted by nonlinear regression to obtain $IC_{50}$ values.

**Supplementary Figure S7**. **Binding of small molecule compounds to ABHD2$^{L33-E425}$.** Binding of small molecule compounds to ABHD2$^{L33-E425}$ was assessed using differential scanning fluorimetry (DSF) following 30 min pre-incubation with 10 μM of the indicated compounds. Dashed lines denote the middle of the melting curve in the fluorescence plot, which corresponds to the peak of the first derivative curve. These lines mark the melting temperature of each sample on the X-axis. The increased melting temperature observed for compound **1, 2, 4**, and **5** were able to stabilise ABHD2$^{L33-E425}$ while no stabilization was observed in the presence of compound **6.**

**Supplementary Figure S8. ABHD2 inhibitors do not inhibit ABHD10 and ABHD11.** ABHD10 and ABHD11 were assayed in a colorimetric assay using p-nitrophenyl butyrate as substrate. 25 nM ABHD11 and 300 nM ABHD10 were incubated for 30 min at 4 °C in the presence of 10 μM compound. Concentrations were 25 nM ABHD11, 300 nM ABHD10, and 500 μM p-nitrophenyl butyrate. Absorbance was monitored at 405 nm for 20 min continuously in 30 sec intervals on a SpectraMax iD3 (MolecularDevices). The unselective ABHD inhibitor MAFP was used as a control.

**Supplementary Figure S9. ABHD2 inhibitors do not affect basal hyperactivation in human sperm.** Paired plots comparing hyperactivation in the presence or absence (DMSO) of ABHD2 inhibitors. Sperm suspensions were treated with either DMSO (vehicle) or 10 μM test compound (**(*R*)-1**, (***R***)**-2**, **4** and **6**). For each of 7 independent experiments, the mean percentage of hyperactive sperm (normalized to motile sperm) across two technical replicates is shown. No statistically significant differences were detected for any comparison (not significant – NS; P > 0.05).

**Supplemental Fig S1**

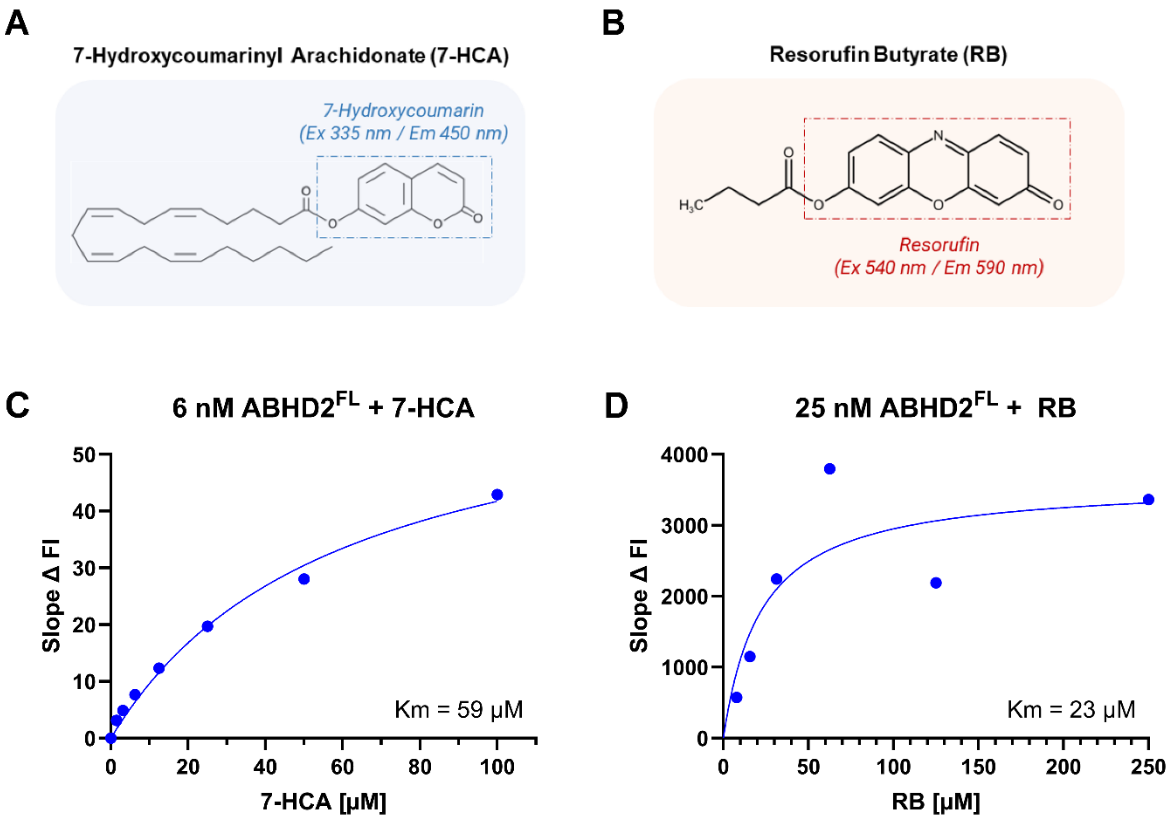

Supplemental Fig S2

A

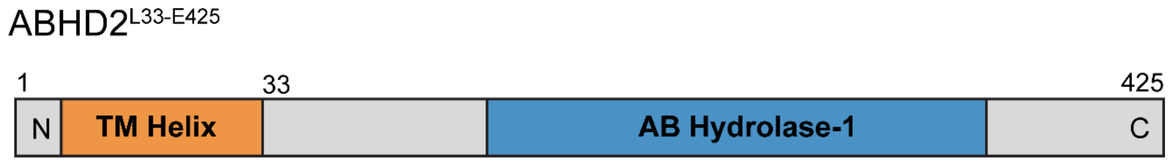


B

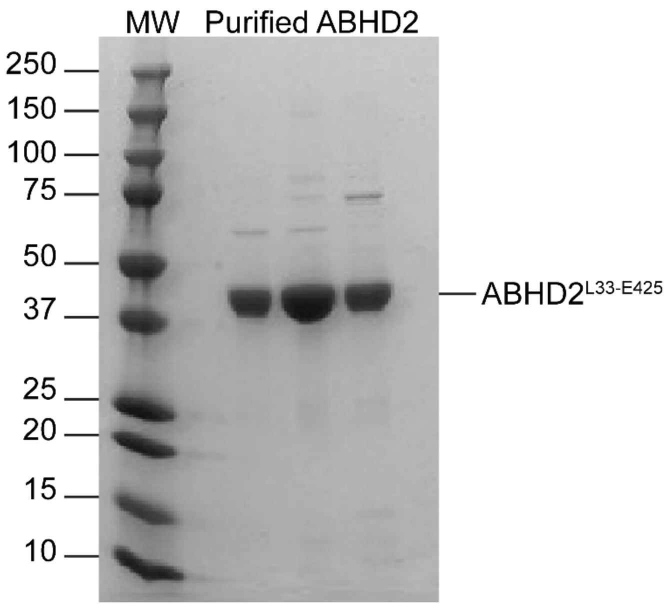


C

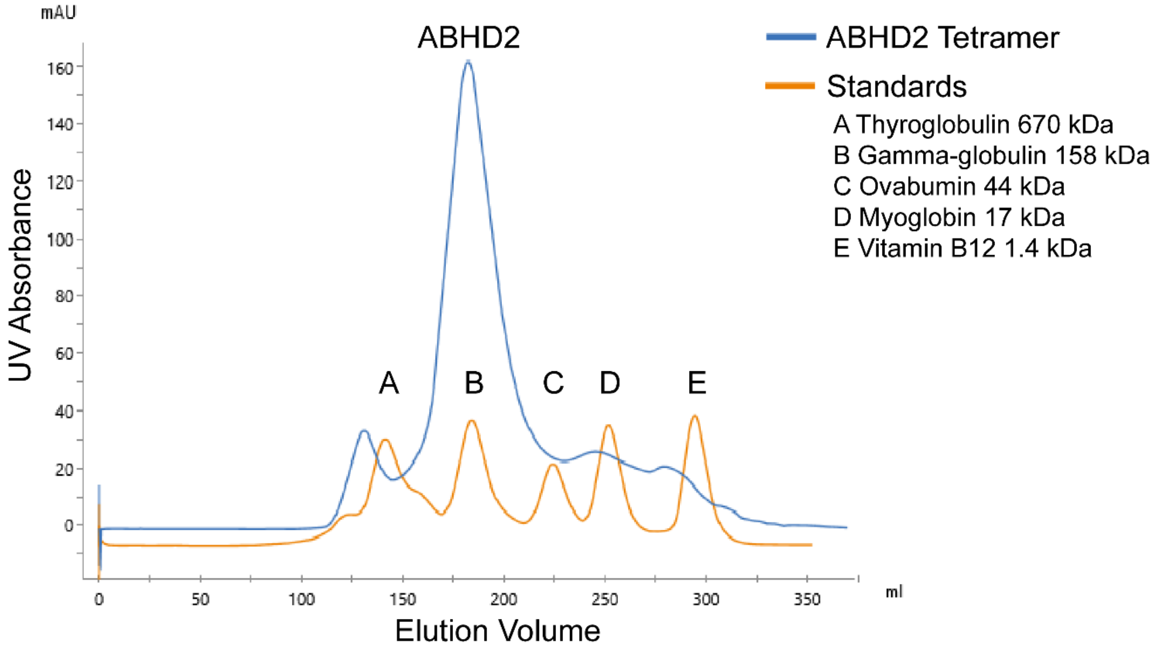

Supplemental Fig S3

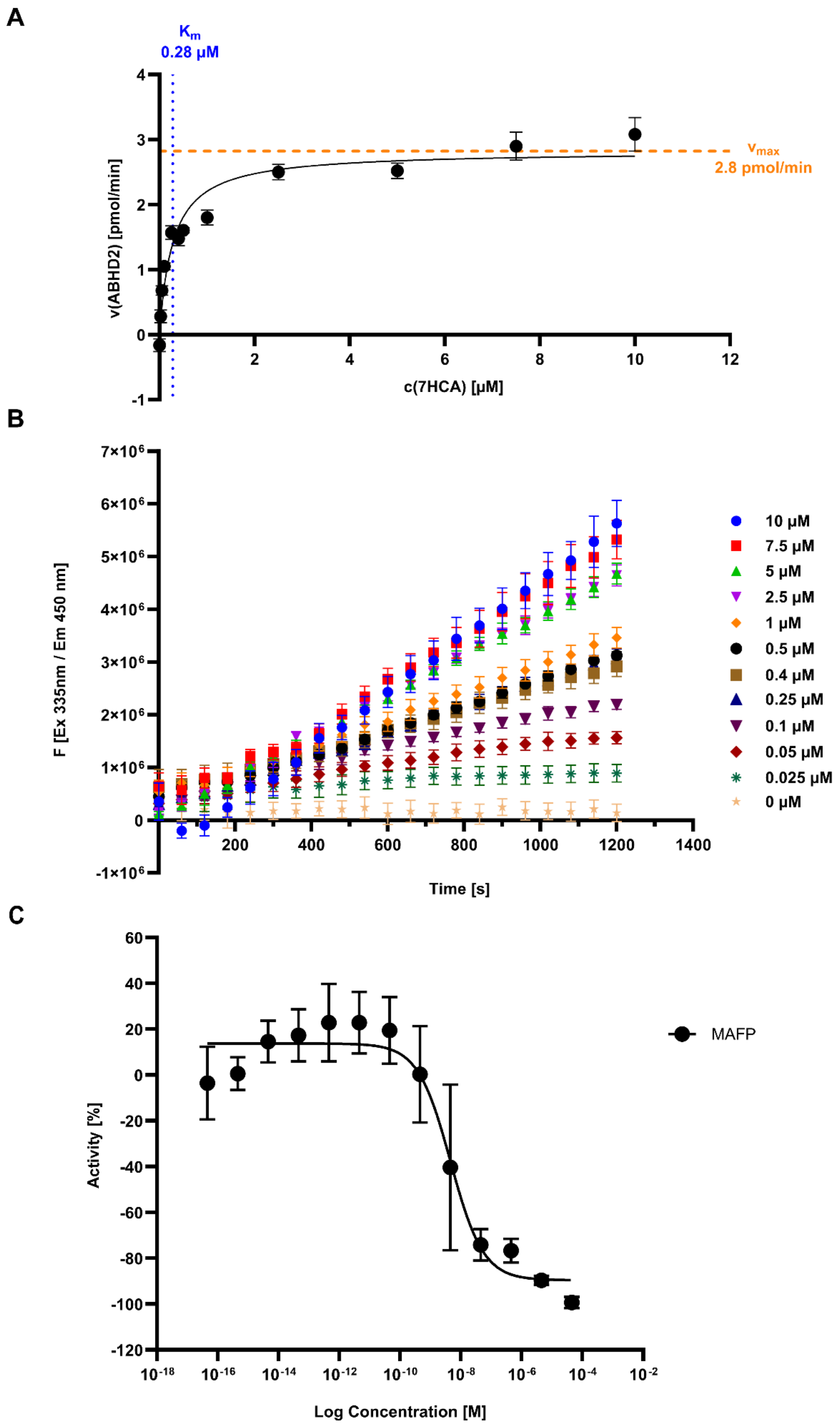

### Supplemental Fig S4

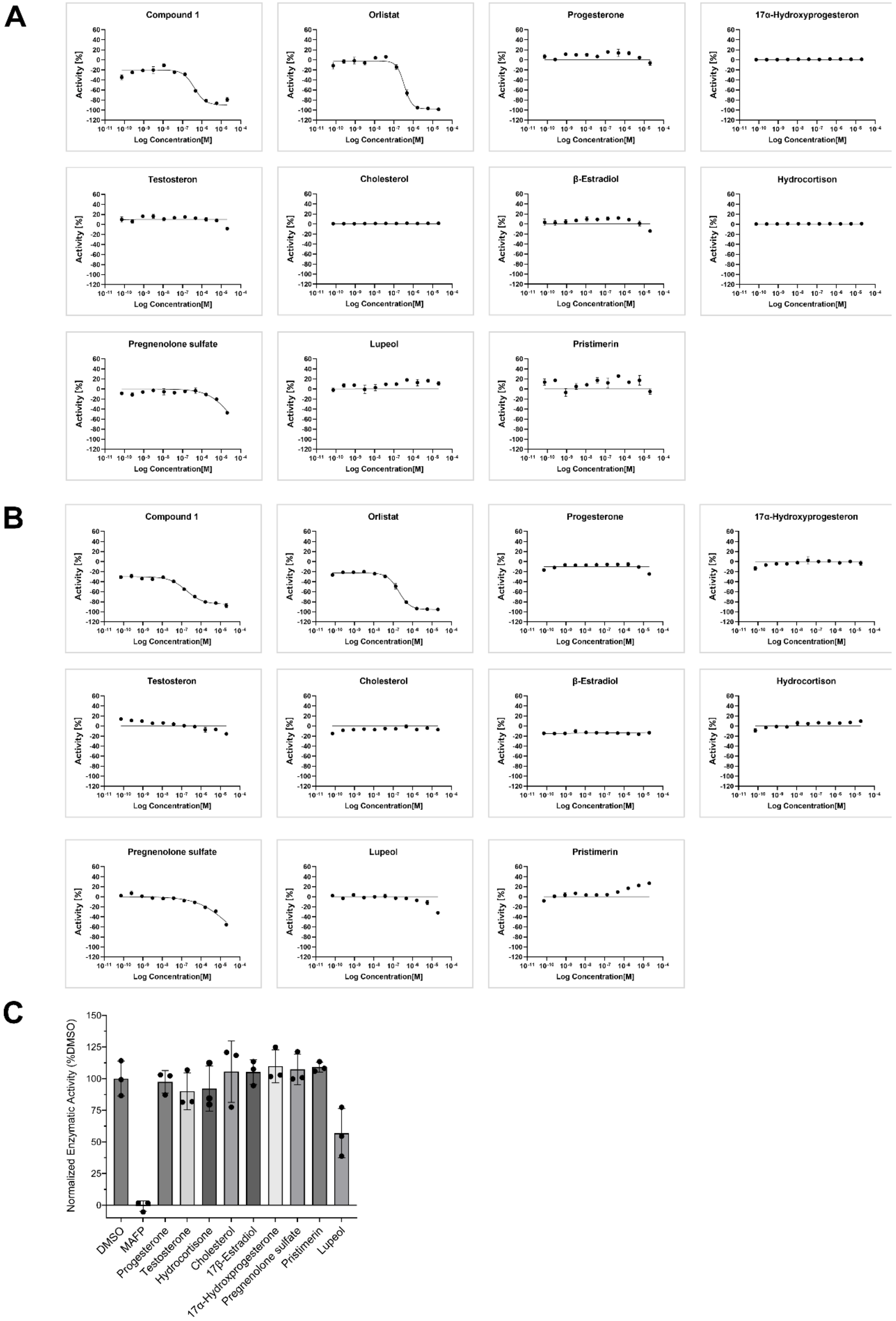

### Supplemental Fig S5

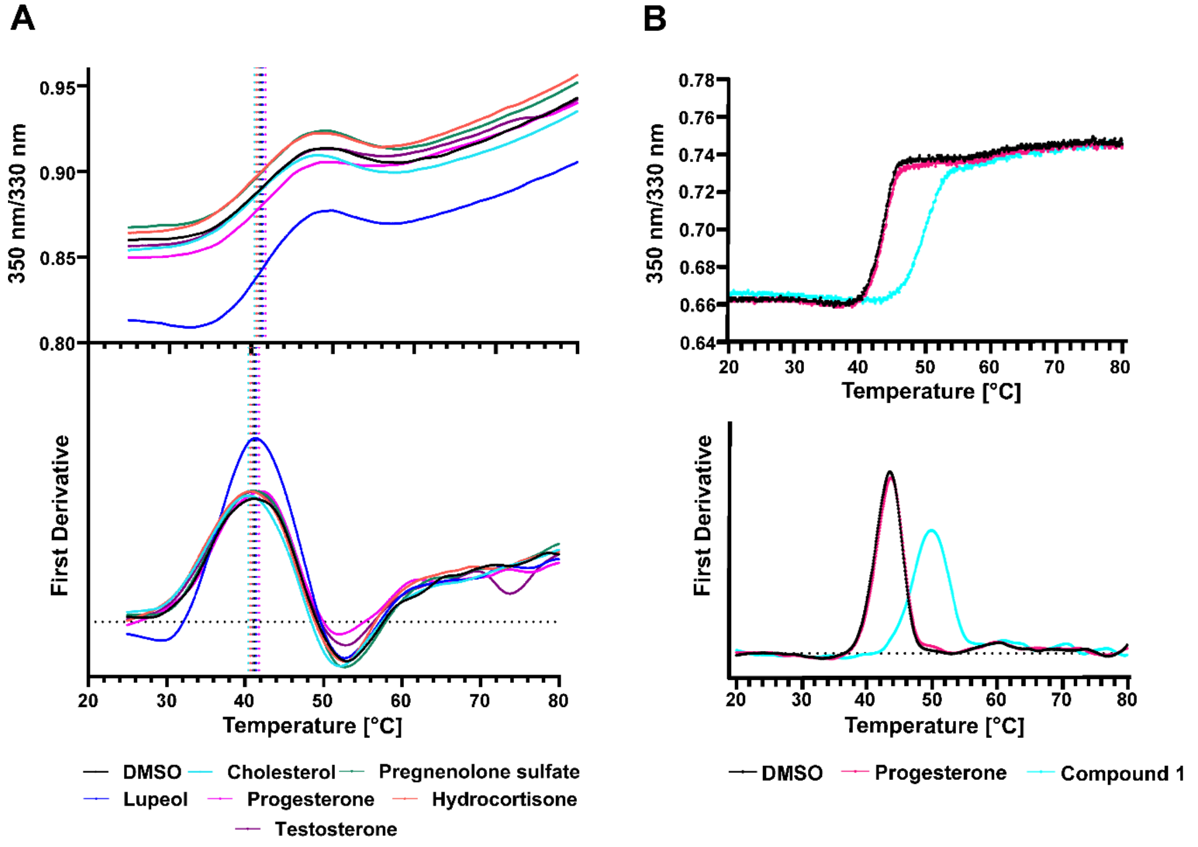


### Supplemental Fig S6

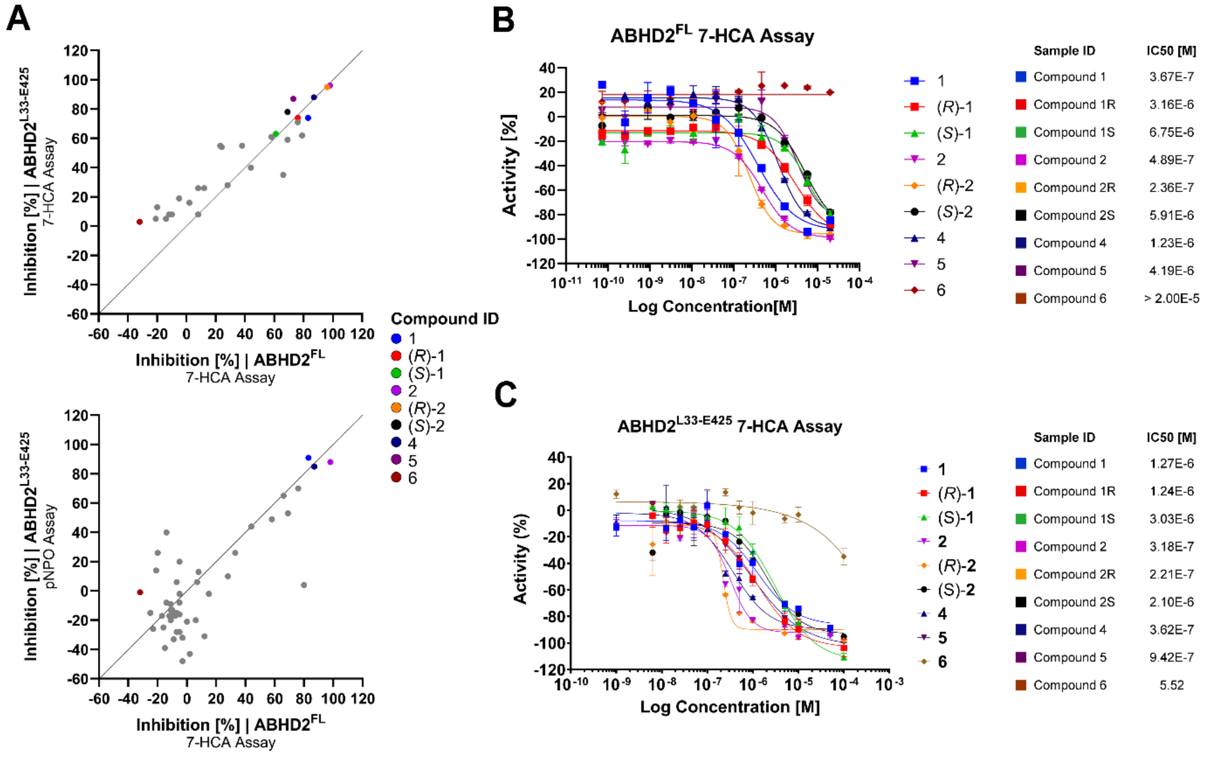

**Supplemental Fig S7**

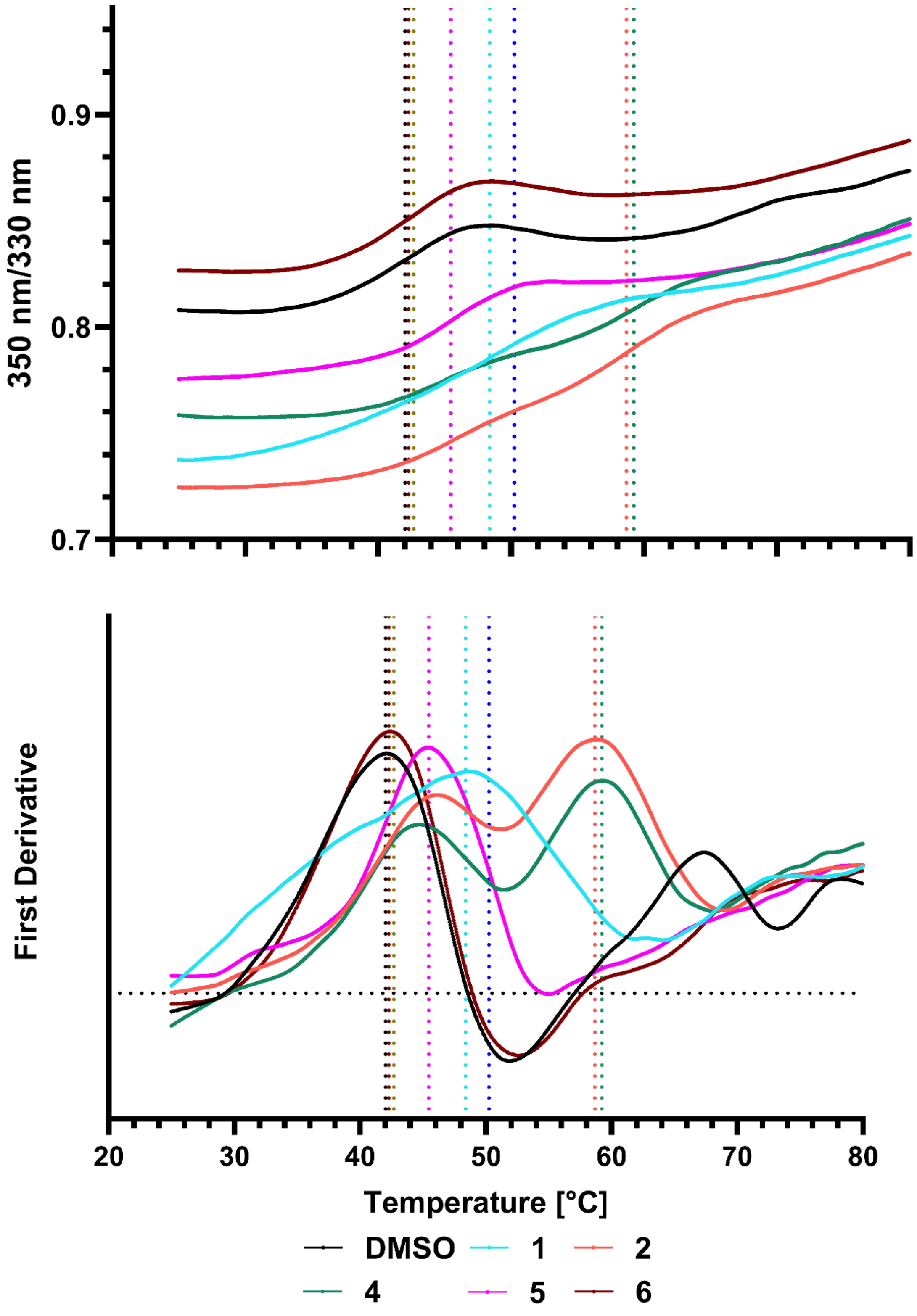

Supplemental Fig S8

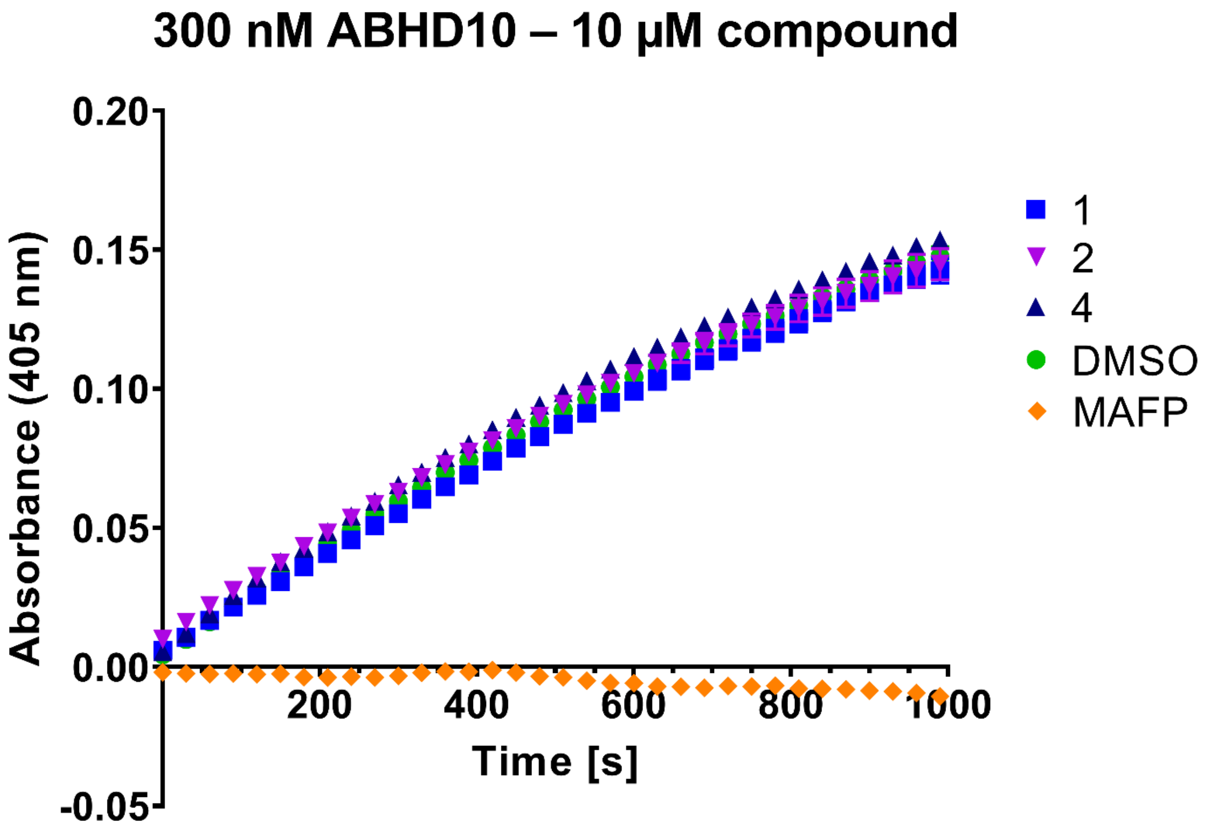


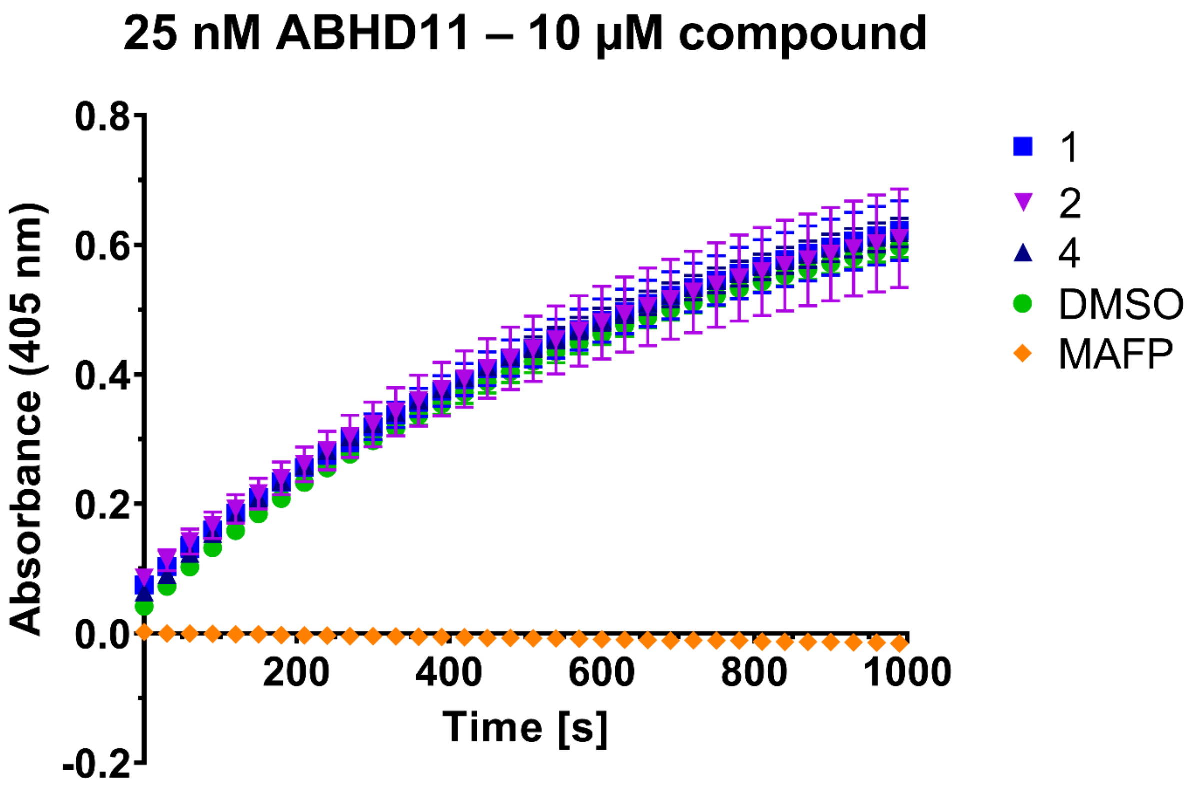

### Supplemental Fig S9

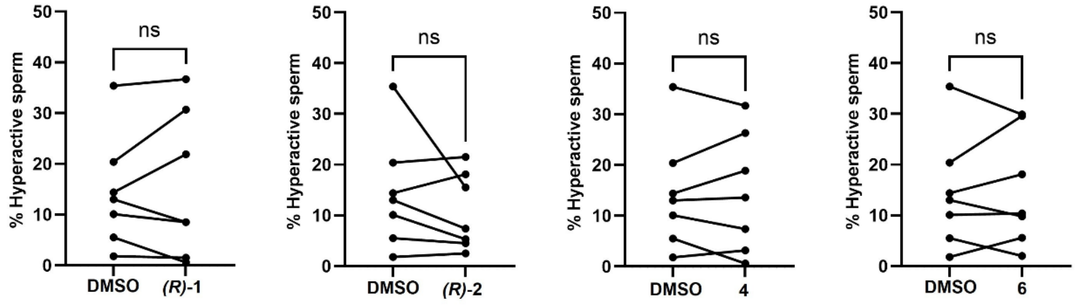
